# AGP-Net: A Universal Network for Gene Expression Prediction of Spatial Transcriptomics

**DOI:** 10.1101/2025.03.09.642276

**Authors:** Yan Yang, Xuesong Li, Liyuan Pan, Guoxun Zhang, Liu Liu, Eric Stone

## Abstract

In the era of high-throughput biology, molecular phenotypes have proven effective in predicting disease states and future trajectories. Transcriptomics, in particular, has enabled the dissection of complex diseases with heterogeneous genetic and environmental aetiology, both aiding diagnosis and augmenting treatment. As improving technology has led to measurements of gene expression at increasing granularity, it has become progressively feasible to resolve disease traits that present locally or with spatial heterogeneity. Principal among these are cancers in which tumour gene expression, while itself heterogenous, exhibits a distinct signature from that of the surrounding tissue. Identifying this signature through molecular phenotyping facilitates specific cancer diagnosis and treatment.

Here, we introduce AGP-Net, a multi-modal foundation framework capable of predicting gene expression from histopathology images. Rather than produce an aggregated estimate of expression for each gene, AGP-Net disaggregates images into spots and attempts to resolve the variation in gene expression across them, thereby providing coarse spatial transcriptomic predictions across the tumour slice and surrounding region. The challenge in doing so is due to data sparsity relative to the dimensionality of the problem: the number of genes and their contextual heterogeneity within and between tissue and cancer types makes it difficult to train a model on the limited data available. The innovation of AGP-Net lies in borrowing strength across similar genes as defined by their textual language descriptions. Our AGP-Net supports datasets with varying gene coverage and facilitates the prediction of gene expression for previously unseen genes based on their textual descriptions. Trained on millions of spots from diverse dataset sources, AGP-Net establishes state-of-the-art performance in zero-shot spatial gene expression prediction, demonstrating its adaptability to generalize across novel scenarios.

## Introduction

Genes encode biological functions that provide insight into both physiology and pathology. Gene expression, the dynamic process by which this encoded information is executed, is particularly useful for understanding the state of a biological sample. In cancer, the most common biological sample is a tumour biopsy, from which gene expression data has proven useful in resolving subtypes and informing treatment decisions. However, the effectiveness of this approach is often limited by the resolution at which gene expression can be measured within the spatial context of a biopsy.

Biological tissues are heterogeneous, and solid tumours are no exception. Bulk gene expression obtained from a tumour sample represents an average across its latent heterogeneity, which may be informative but disregards any variability present in the tumour [1]. The spatial resolution of this variability across the tumour sample allows for more refined diagnoses and treatments. In particular, it becomes easier to: i) identify the (sub)type of cancer present and potential mosaicism; ii) define the margin distinguishing the tumour from normal tissue; iii) quantify the degree of cancer progression [2].

In recent years, spatial transcriptomics has emerged as a means of obtaining in situ measurements of local gene expression within a tissue of interest. By quantifying gene expression across an array of spots, spatial transcriptomics provides spatially resolved gene expression data, enabling deep insights into biological functions and offering crucial support for clinical diagnostics [3]. However, the widespread adoption of spatial transcriptomics is constrained by the high experimental costs associated with techniques such as sequencing [4]. These limitations have driven the development of computational pathology methods that infer spatial transcriptomics directly from hematoxylin and eosin (H&E)-stained histopathology images, offering a cost-effective and scalable alternative [5–10].

Existing methods, however, face considerable challenges in generalizability, primarily due to limited dataset availability and variability in data acquisition platforms. For example, the STNet dataset [5] consists of only 68 histopathology images from 23 patients, while popular platforms like 10x Visium [11] and Stereo-seq [12] differ substantially in spatial resolution and gene coverage [13]. These limitations confine models to being trained and tested on samples of histopathology images collected under similar conditions and restrict their applicability to supervised learning tasks. In particular, the majority of them can only predict the expression of genes observed when training the model. Such constraints significantly hinder the broader applicability of these models across diverse clinical and research scenarios.

Although recent efforts have focused on curating spatial transcriptomics datasets from various published studies [14, 15], the scope and breadth of these datasets remain insufficient to develop a robust foundational model. Such a model would require extensive data diversity to be generalized across the wide range of environments, experimental protocols, and clinical settings encountered in real-world applications [16]. Consequently, this limitation hinders the broader adoption of spatial transcriptomics for advancing precision medicine and clinical diagnostics.

To overcome the challenges, we reframe the gene expression prediction task as a multi-modal learning problem that links textual descriptions of genes with histopathology images. This results in <u>a</u> universal <u>g</u>ene expression <u>p</u>rediction <u>net</u>work (AGP-Net). It (Fig. 1a) is trained on a large-scale dataset curated from multiple sources (Fig. 1b-c). By leveraging textual language descriptions, AGP-Net predicts gene expression through correlations between histopathology images and the textual annotations of the genes of interest. This approach not only accommodates datasets with varying gene coverage but also enables the prediction of gene expression for unseen genes based on their textual descriptions. To fully unleash the model performance across diverse scenarios, we combine existing datasets with curated resources designed to enhance data diversity and scale. Specifically, histopathology images paired with textual captions can be leveraged to construct datasets representing artificial genes with binary expression. In this framework, the textual caption acts as a proxy for an artificial gene: samples paired with a caption are labeled as having positive expression, while unpaired ones are labeled as having zero expression. Each image is treated as a single spatial annotation point, symbolifying the representation of gene expression across the entire image. This setup provides coarse supervision signals for training the foundational model. When combined with real spatial transcriptomics datasets, this strategy enables AGP-Net to deliver exceptional performance across a wide range of clinical tasks, positioning it as a versatile foundational model for spatial transcriptomics. We validate the clinical and technical significance of AGP-Net across various applications. Computationally, AGP-Net directly predicts gene expression from histopathology images obtained under varying conditions and environments, while also enabling effective gene expression imputation. Operationally, it supports advanced clinical tasks such as tumor detection and cancer subtyping, demonstrating robust performance across histopathology images collected from diverse settings.

**Fig. 1:**
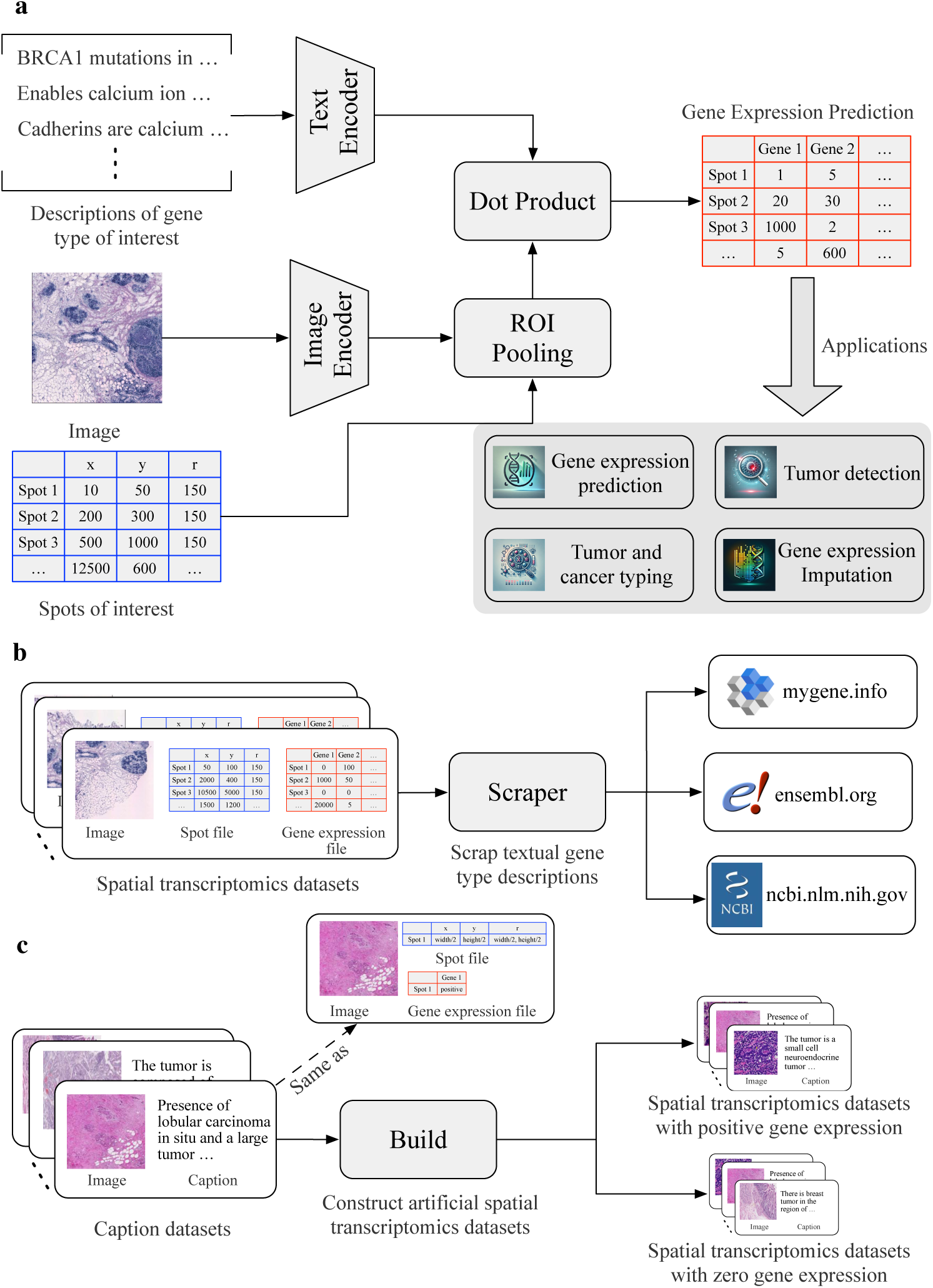
Framework overview. **a**, the proposed method consists of a text encoder and an mage encoder. Text descriptions of the gene of interest and histopathology images are encoded separately using the text and image encoders. To predict gene expression for specific spots of interest, ROI pooling is applied to the image encoder outputs based on a spot file that defines the regions corresponding to each spot. Gene expression predictions are calculated as the dot product between the outputs of the text encoder and the ROI pooling results. This framework supports various applications, such as tumor detection and gene expression imputation. **b**, to train the method, we perform web scraping to retrieve textual descriptions of genes from diverse sources for using with spatial transcriptomics datasets. **c**, the caption dataset of histopathology images s used to create artificial spatial transcriptomics datasets, by considering the paired histopathology image and textual descriptions with positive gene expression, and non-paired ones with zero gene expression.

## Result

### Gene expression prediction

By leveraging free-form textual descriptions of genes, AGP-Net allows the prediction of expression for a wide range of genes, irrespective of whether those genes are seen during the training phase. Feeding a whole histopathology image, language descriptions of target genes, and spots of interest as inputs, AGP-Net predicts log-transformed expression values, without any further fine-tuning. We evaluate AGP-Net on a diverse set of spatial transcriptomics datasets comprising 13 dataset collections, spanning multiple experimental platforms and clinical conditions. For each source, the 250 most highly expressed genes are selected based on expression profiles, following the protocol in [5]. Model performance is quantified using the Pearson correlation coefficient (PCC) between predicted and ground truth gene expression values, with the first quantile (PCC@F), the mean (PCC@M), and the third quantile (PCC@S) of PCC reported across all target genes, as shown in Tab. 1.

**Table 1:** Gene expression prediction performance of AGP-Net on 13 spatial transcriptomics dataset collections.

| Dataset | PCC@F | PCC@M | PCC@S |
| --- | --- | --- | --- |
| Subudhi *et al.* [17] | 0.373 | 0.415 | 0.477 |
| Mitamura *et al.* [18] | 0.302 | 0.325 | 0.391 |
| Ji *et al.* [19] | 0.494 | 0.530 | 0.580 |
| Sans *et al.* [20] | 0.153 | 0.184 | 0.231 |
| Cortese *et al.* [21] | 0.064 | 0.202 | 0.356 |
| Yu *et al.* [22] | 0.161 | 0.197 | 0.239 |
| Lyubetskaya *et al.* [23] | 0.087 | 0.170 | 0.289 |
| Zou *et al.* [24] | 0.159 | 0.206 | 0.283 |
| Erickson *et al.* [25] | 0.341 | 0.359 | 0.397 |
| Andersson *et al.* [26] | 0.277 | 0.329 | 0.396 |
| Coutant *et al.* [27] | 0.158 | 0.231 | 0.314 |
| Kuppe *et al.* [28] | 0.086 | 0.175 | 0.276 |
| Maynard *et al.* [29] | 0.177 | 0.211 | 0.246 |

AGP-Net demonstrates robust generalization and adaptability across diverse clinical and experimental conditions, highlighting its potential in advancing precision medicine. For instance, on the Ji *et al.* [19] dataset, AGP-Net achieves a mean Pearson correlation coefficient (PCC@M) of 0.530, underscoring its ability to accurately predict gene expression profiles. The accurate predictions, particularly of cancer-related biomarkers, hold significant value in aiding diagnostics and guiding personalized treatment strategies. To further showcase the utility of AGP-Net, we provide examples of spatial gene expression predictions compared to ground truth values for key cancer-related markers in Fig. 2. From left to right of Fig. 2, we respectively show the histopathology image, spatial distribution of ground truth gene expression, spatial distribution of predicted gene expression, and the distribution comparison between the ground truth and predictions. The example genes are KRT5 (Fig. 2a) and SPAR (Fig. 2b) for skin cancer, as well as FAN (Fig. 2c) and ERBB (Fig. 2d) for breast cancer. The results demonstrate an excellent visual correspondence between predicted and ground truth gene expression, even in challenging out-of-distribution scenarios (i. e., cross-dataset evaluations). The robustness and versatility highlight the capability of AGP-Net to capture gene expression variations across a wide range of clinical and experimental settings, making it a valuable tool for translational biomedical research and clinical applications.

**Fig. 2:**
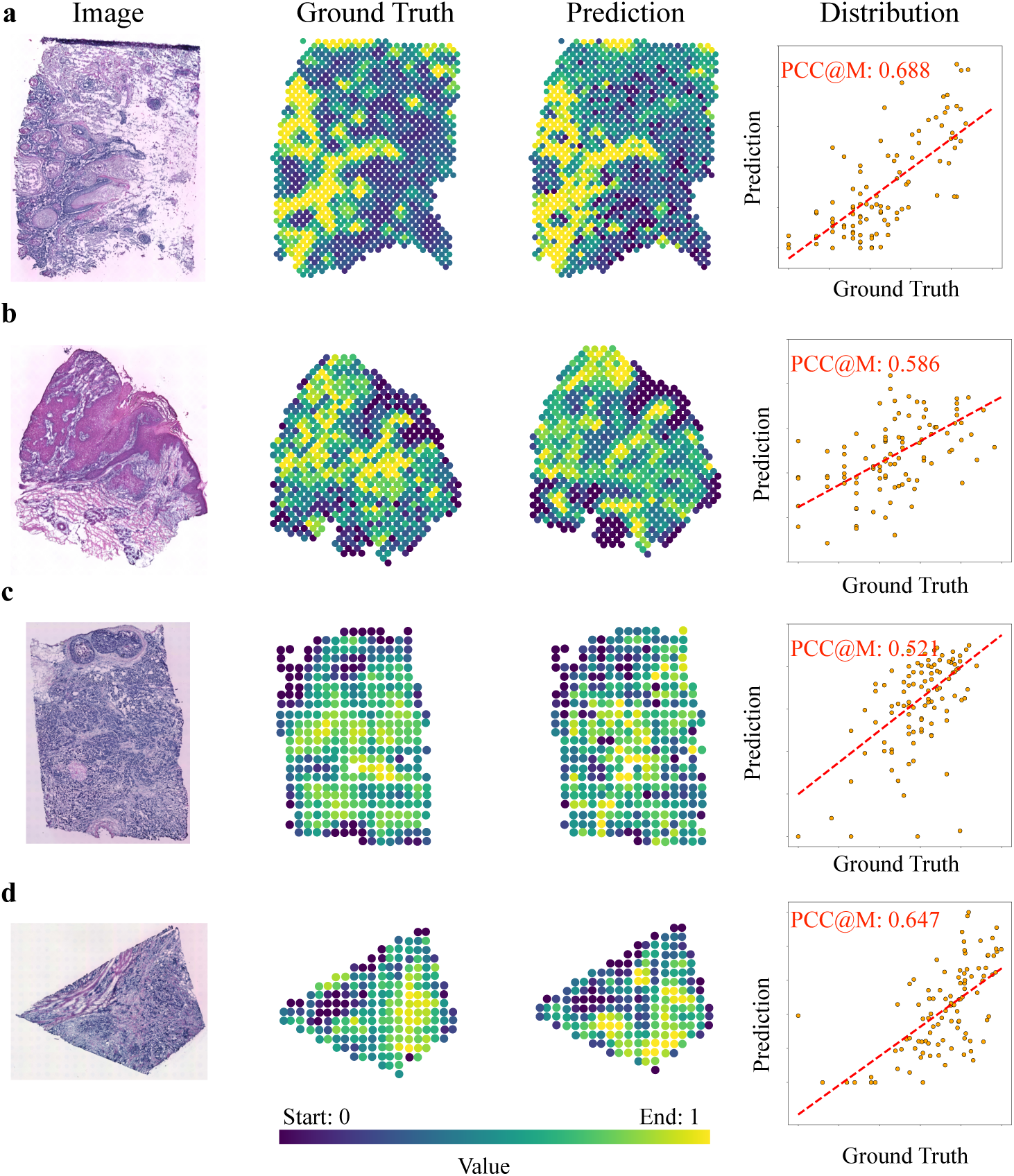
Spatial gene expression prediction. From the left to right, we show the histopathology image, ground truth expression, predicted gene expression, and detail distribution comparisons between prediction and ground truth gene expression. For the distribution comparisons, we sample 100 spots for visualization clarity, and the red dotted line represents the regression line between the prediction and ground truth. **a** and **b**, samples from Ji *et al.* [19] dataset respectively explores the skin diseases related genes of KRT5 and SPARC. **c** and **d**, samples from Andersson *et al.* [26] datasets respectively explores genes of FAN and ERBB2 that are correlating with breast diseases.

### Tumor Detection

Our method precisely aligns histopathology images with spatial gene expression data by linking textual descriptions of genes to histopathology images. This approach enables a CLIP-like zero-shot recognition^1^ [30] capability in the biomedical context. A direct and specific application of this method is the detection of tumor presence, achieved by analyzing the expression levels of tumor-related markers or markers of healthy/benign tissue. We demonstrate the advantages of AGP-Net over previous state-of-the-art zero-shot frameworks, as shown in Fig. 3.

**Fig. 3:**
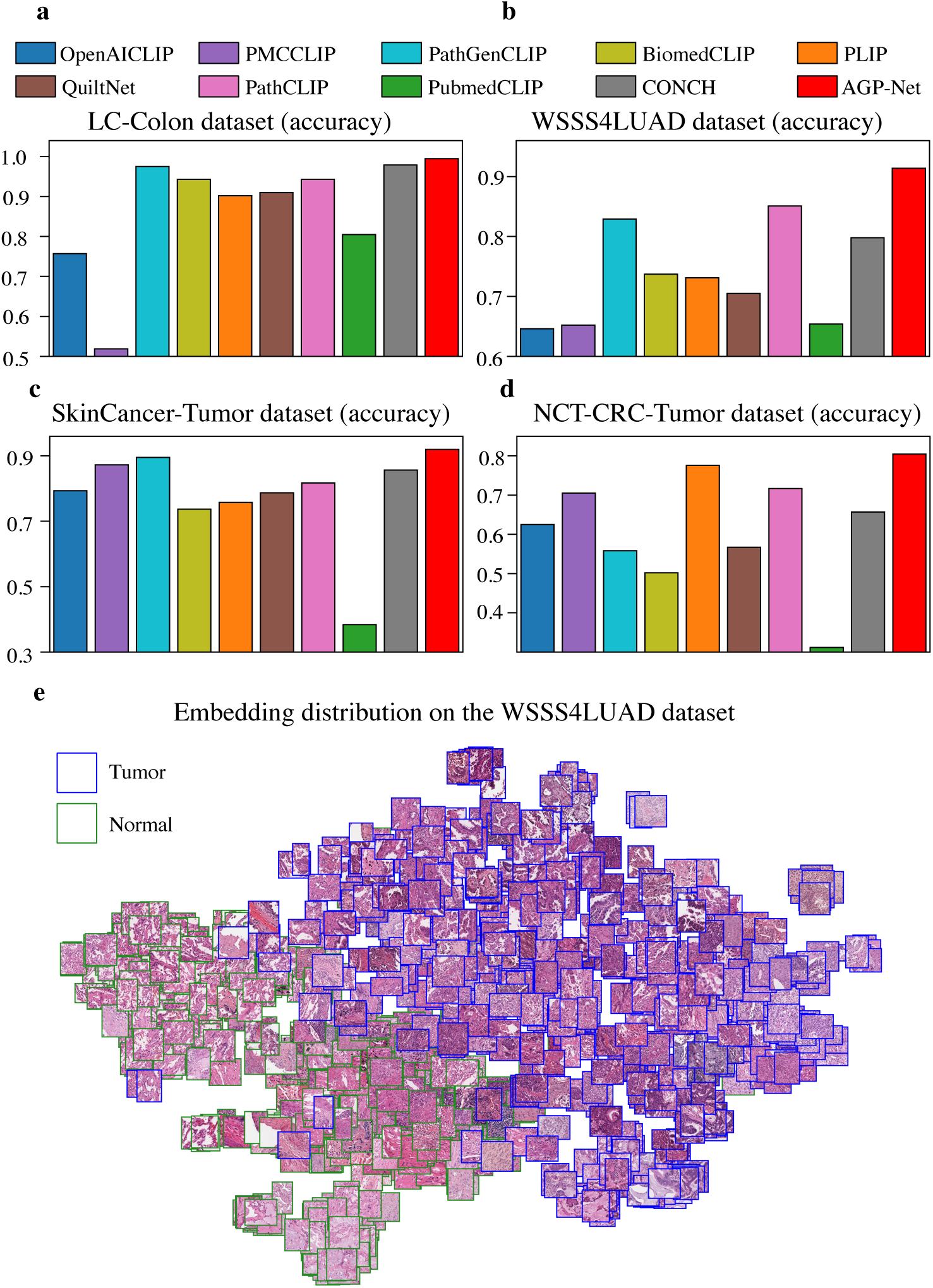
Comparison of tumor detection. **a**-**d**, the accuracy of state-of-the-art zero-shot methods on the LC-Colon dataset [31], WSSS4LUAD dataset [32], SKinCancer-Tumor dataset [33], and NCT-CRC-Tumor dataset [34]. We compare with OpenAICLIP [30], PMCCLIP [50], PathGenCLIP [44], BiomedCLIP [51], PLIP [46], QuiltNet [47], PathCLIP [48], PubmedCLIP [52], and CONCH [16], coloring each of them into distinct colors. **e**, distribution of 1000 sampled embeddings of histopathology images from the WSSS4LUAD dataset. The dimension reduction is performed with t-SNE [53]. We highlight the tumor and normal tissue respectively in blue and forest green.

By incorporating textual descriptions of genes of interest, AGP-Net allows querying both actual genes and artificial genes (as indicated above, *e.g.*, using terms such as ‘tumor’ directly). This functionality facilitates direct comparisons between the expression patterns of artificial genes like ‘tumor’ and ‘normal’ tissue for tumor detection tasks. To evaluate this capability, we conduct tumor detection experiments across four datasets: the LC-Colon dataset [31] (Fig. 3a), WSSS4LUAD dataset [32] (Fig. 3b), SKinCancer-Tumor dataset (Fig. 3c), and NCT-CRC-Tumor dataset (Fig. 3d). These datasets encompass both scenarios: some contain separate slides for tumor and normal tissues, while others include both within the same slide. The latter two datasets are variations of the SKinCancer dataset [33] and NCT-CRC dataset [34], respectively, with their classes consolidated into two categories: tumor and normal tissues. Model performance is evaluated using accuracy.

AGP-Net consistently outperforms previous state-of-the-art methods across all evaluations. For example, on the WSSS4LUAD dataset, AGP-Net achieves an accuracy of 0.914, substantially surpassing the next best method, which achieves 0.851. As illustrated in Fig. 3e, the feature distributions of histopathology images from the WSSS4LUAD dataset [31] demonstrate the method’s effectiveness in distinctly separating tumor regions (highlighted by blue rectangles) from normal tissues (highlighted by forest green rectangles). This clear segregation underscores the superior performance of AGP-Net and its ability to accurately capture and utilize underlying patterns in histopathology images for reliable tumor detection.

### Tumor and Cancer Typing

Expanding on capabilities of AGP-Net in tumor detection, it demonstrates robust zero-shot tumor and cancer typing by leveraging spatial gene expression patterns. Performance comparisons with state-of-the-art methods are presented in Fig. 4 across six dataset collections. The BACH dataset (Fig. 4a) [35] classifies tissue into four categories: benign, in-situ carcinoma, invasive carcinoma, and normal. The BreakHis dataset [36](Fig. 4b) distinguishes between benign and malignant tumors. Both the LC-Lung [31] (Fig. 4c) and LungHist700 [37] (Fig. 4d) datasets categorize samples into benign tissue, adenocarcinoma, and squamous cell carcinoma. The SICAP dataset [38] (Fig. 4e) differentiates non-cancerous tissue from Gleason grades 3-5 [39]. Lastly, the SkinCancer dataset [33] (Fig. 4f) examines 12 anatomical compartments and 4 types of neoplasms. Similarly, accuracy is reported as the evaluation metric.

**Fig. 4:**
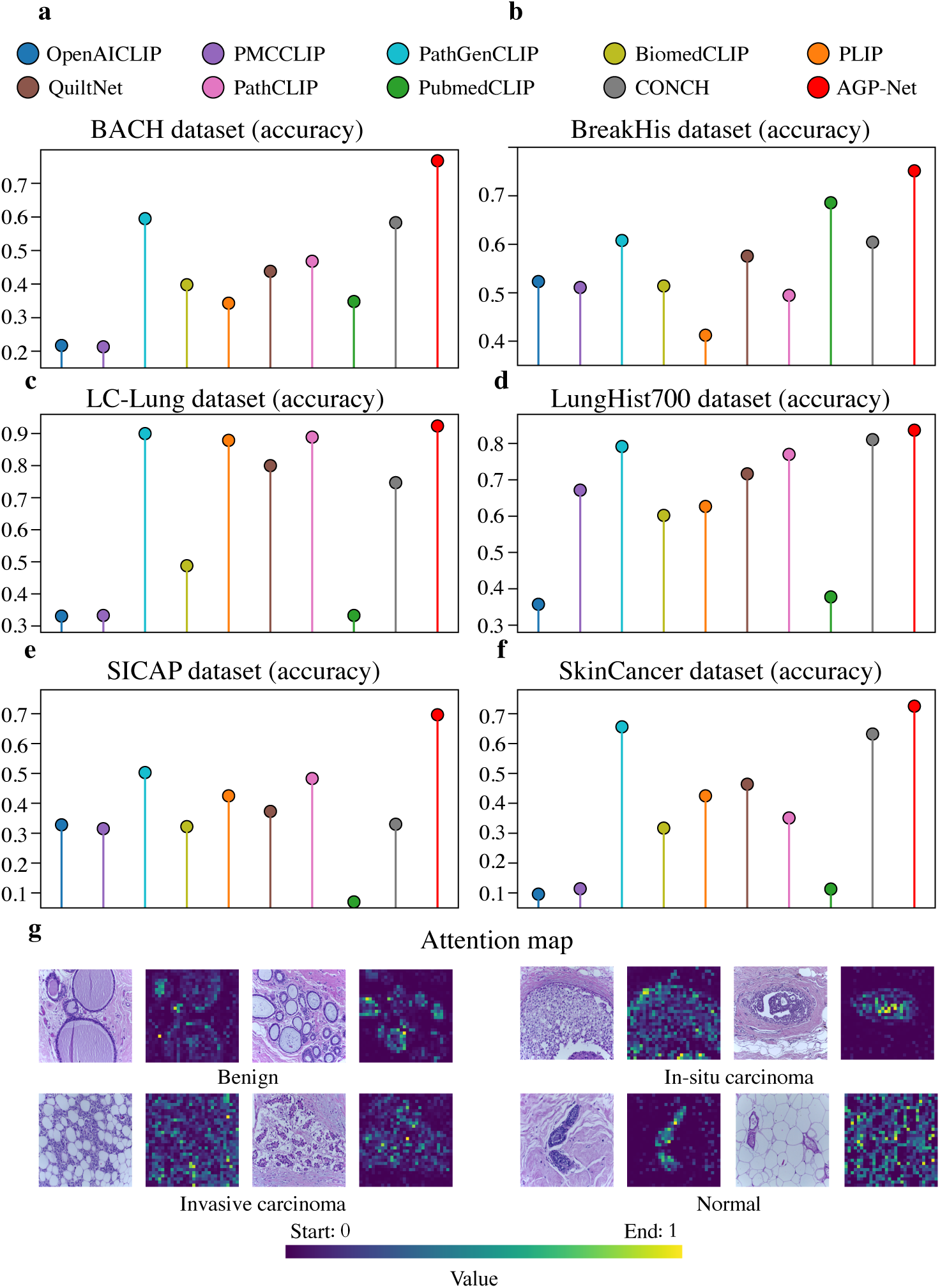
Comparison of tumor and cancer typing. **a**-**f**, the accuracy of state-of-the-art zero-shot methods on the BACH dataset [35], BreakHis dataset [36], LC-Lung [31], LungHist700 [37], SICAP dataset [38], and SkinCancer dataset [33]. We compare with OpenAICLIP [30], PMCCLIP [50], PathGenCLIP [44], BiomedCLIP [51], PLIP [46], QuiltNet [47], PathCLIP [48], PubmedCLIP [52], and CONCH [16], coloring each of them into distinct colors. **g**, attention maps of our AGP-Net for histopathology images from the BACH dataset. The warmer the color, the more discriminative the region.

AGP-Net consistently demonstrates superior performance in tumor and cancer typing, significantly surpassing previous zero-shot methods. For example, when classifying tumor progression on the BACH dataset, AGP-Net achieves an accuracy improvement of 0.173 over the next-best method. Similarly, for grading cancers on the SICAP dataset, AGP-Net achieves an accuracy improvement of 0.193. These results highlight the advanced ability of the model to capture subtle spatial gene expression patterns. As shown in Fig. 4g, the attention maps generated by AGP-Net for histopathology images in the BACH dataset dynamically focus on discriminative regions, providing interpretability and insights into its decision-making process.

### Gene Expression Imputation

The capability of AGP-Net extends beyond zero-shot gene expression prediction to effective imputation of gene expression values, demonstrating its potential to enable robust spatial transcriptomics profiling. For gene expression imputation, a subset of spatial spots and their corresponding gene expression data is provided as supervised training data. AGP-Net leverages this sparse information to infer the expression levels of genes at other spatial spots within the same slide. This imputation ability is critical for applications where acquiring complete gene expression profiles is costly, time-intensive, or technically challenging.

The comparison with state-of-the-art methods is illustrated in Fig. 5. We bench-marked AGP-Net against representative zero-shot histopathology frameworks on four high-quality datasets: the Ji *et al.* [19] dataset (Fig. 5a), Andersson *et al.* [26] dataset (Fig. 5b), Maynard *et al.* [29] dataset (Fig. 5c), and Erickson *et al.* [25] dataset (Fig. 5d). For each method, a linear layer is attached to the image encoder to perform the imputation task. To account for the variability of the imputation tasks, five runs are performed for each method on each slide image of the datasets. The PCC@M (i. e., mean Pearson correlation coefficient) is used as the evaluation metric.

**Fig. 5:**
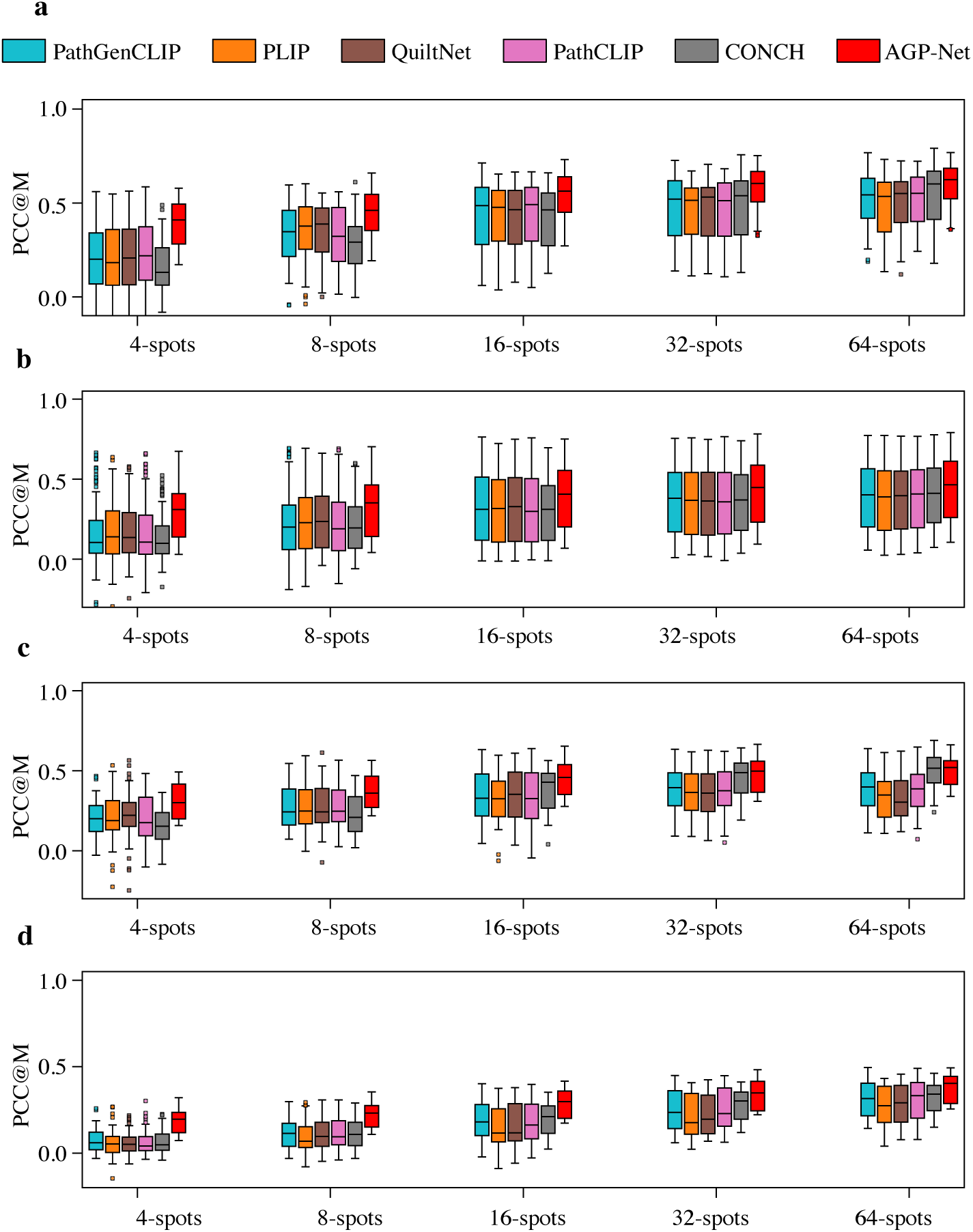
Comparison of gene expression imputation. **a**-**d**, the PCC@M (i. e., mean Pearson correlation coefficient) on the Ji *et al.* [19] dataset, Andersson *et al.* [26] dataset, Maynard *et al.* [29] dataset, and Erickson *et al.* [25] dataset. The x-axis represents the number of spots used for training in histopathology images, starting at 4 and doubling with each subsequent step. To account for the variability of the imputation tasks, 5 runs are performed for each method on each slide image of the datasets. We note the state-of-the-art methods are unable to effectively predict gene expression when training when less than 4 spots. Thereby, we start the comparisons with 4 spots.

AGP-Net consistently outperformed competing methods in gene expression imputation across all benchmarks. Its superior performance is particularly evident in scenarios with limited spatial spots. For example, on the Maynard *et al.* [29] dataset, AGP-Net achieved a PCC@M improvement exceeding 0.2 over the next-best method. These results highlight AGP-Net’s robustness across diverse datasets and its precision in imputing gene expression patterns, even in data-sparse settings. Its ability to reconstruct gene expression landscapes enables detailed exploration of tissue microenvironments, a deeper understanding of disease mechanisms, and the development of targeted therapeutic interventions, making it a valuable tool for advancing spatial biology and biomedical research.

## Discussion

In this study, we present AGP-Net, a foundation model designed for spatial transcriptomics and histopathology images, specifically targeting zero-shot gene expression prediction. By framing the genes of interest using free-form textual descriptions, AGP-Net enables flexible and scalable gene expression prediction across a variety of datasets without requiring task-specific retraining. This capability makes it well-suited for applications where new gene markers or disease-specific targets are introduced, as it seamlessly integrates textual queries to guide prediction.

The strength of AGP-Net lies in its ability to align spatial transcriptomics data with histopathology images through a shared latent space. This alignment not only facilitates gene expression prediction but also enables downstream tasks such as understanding the spatial organization of gene activity and gene expression imputation. Fundamentally, AGP-Net represents a promising solution for zero-shot gene expression prediction, enabling efficient clinical diagnosis under a wide range of contexts. Furthermore, AGP-Net consistently outperforms state-of-the-art methods in both tumor detection and cancer typing tasks across multiple dataset collections. For example, in tumor progression classification on the BACH dataset, AGP-Net achieves significant accuracy improvements over prior methods.

Through the analysis of feature distributions and attention heatmaps produced by AGP-Net, the model interpretability is effectively demonstrated. These visualizations showcase its ability to group histopathology images with distinct features into well-defined clusters and to pinpoint discriminative regions that are potentially biologically and diagnostically significant. By highlighting these regions within histopathology images, AGP-Net offers valuable insights into its decision-making process. The attention maps, in particular, emphasize key areas associated with discriminative features, enabling a deeper understanding of the underlying biological processes. This interpretability is a crucial strength of AGP-Net, enhancing its reliability and fostering greater confidence in its use for clinical applications.

Despite its promising capabilities, AGP-Net has certain limitations that warrant further exploration. One notable limitation is its reliance on high-quality spatial transcriptomics and histopathology datasets for training and evaluation. Extending its applicability to accommodate variations in staining techniques, imaging protocols, and batch effects could significantly enhance its robustness and reliability for real-world clinical applications. Additionally, incorporating domain-specific knowledge, such as curated gene ontologies or molecular pathway information, could refine its predictions and broaden its usability across diverse biomedical contexts. Another avenue for future work is the exploration of AGP-Net in multi-modal analysis by integrating additional data types, such as imaging mass spectrometry or single-cell RNA sequencing data, to further enhance its predictive power.

In conclusion, AGP-Net represents a significant advancement in bridging spatial transcriptomics and histopathology images for zero-shot gene expression prediction and related tasks. Its ability to generalize across datasets, interpret biological processes, and deliver reliable results highlights its potential as a transformative tool in both research and clinical domains. Future efforts will focus on addressing the aforementioned limitations, expanding its applications to other modalities, and integrating it into clinical workflows to realize its full potential in personalized medicine.

## Methods

### Network Architecture of AGP-Net

The core architecture of AGP-Net employs a dual-encoder design, comprising a visual image encoder and a text encoder. The image encoder is based on the ViT-B/16 transformer backbone [30], enhanced with interleaved convolutional layers inserted before every transformer block. Specifically, for the convoltion layers, we use the conditional positional embedding block from [40]. Within each transformer block, the attention mechanism operates on a window size of 224 × 224 pixels, and the interleaved convolutional layers improve information flow across different windows of the histopathology images, enabling the encoder to transform each image into a comprehensive feature map. The text encoder utilizes a transformer-based architecture adapted from [30]. It is designed to handle textual inputs of up to 256 tokens and encodes each description into a 512-dimensional feature vector. For text processing, the encoder employs a byte pair encoding tokenizer [41] for tokenizations.

To predict gene expression at a specific spot within a histopathology image, whether derived from real or artificial spatial transcriptomics datasets, the feature map generated by the visual encoder is pooled using the roi align function [42] according the spots of interest, producing a 512-dimensional feature vector corresponding to the region of interest. This visual feature vector is then combined with the 512-dimensional feature vector generated by the text encoder through a dot product operation to estimate gene expression. Both feature vectors are *ℓ*_2_-normalized prior to the dot product. To accommodate real and artificial spatial transcriptomics datasets, two learnable scale parameters are used to adjust the dot product output, mapping it to continuous gene expression values for real datasets and more binary-like gene expression values for artificial datasets.

### Dataset

For the real spatial transcriptomics datasets, we focus on data derived from human tissue. The datasets are sourced using the platforms described in [14, 15], which aggregate publicly available spatial transcriptomics data. A subset of these datasets is allocated for training, while the remaining portion is designated for evaluating the gene expression prediction performance (see Tab. 1 for details on the testing data). To optimize the training process, we preprocess the data by selecting the 250 most highly expressed genes for each histopathology image. The images are padded to ensure that the dimensions are divisible by 1792, then tiled into patches of 1792 × 1792 resolution. Patches with fewer than 32 spatial spots or fewer than 64 highly expressed genes (as determined by [43]) are excluded. This preprocessing results in 11K patches, each with a resolution of 1792 × 1792, comprising 1.4M spatial spots and 30K distinct genes. To generate natural language descriptions for the selected genes, we perform web scraping using three sources, which are mygene.info, ensembl.org, and ncbi.nlm.gov. For the artificial spatial transcriptomics datasets, we incorporate data from [44–48], creating a combined dataset of 2.7M histopathology image patch and caption pairs. This diverse and high-quality artificial dataset further enhances the training efficacy of the model by providing robust and varied training examples.

### Model Training

We implement AGP-Net using the *PyTorch* framework [49]. The network is trained for 30 epochs jointly on the real and artificial spatial transcriptomics datasets. During training, each batch is controlled to contain at least 256 spots from the real transcriptomics dataset, corresponding to 8 patches of resolution 1792 × 1792, and 768 spots from the artificial dataset. For the real transcriptomics dataset, we minimize the following loss function: 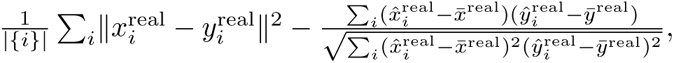 where |{*i*}| denotes the batch size, and the summation operator 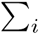 enumerates over the predicted gene expression 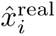 and log-transformed ground truth gene expression 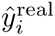 within the batch. Here, 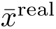 and 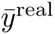 represent the mean of the predicted and ground truth gene expressions for the batch, respectively. For the artificial transcriptomics dataset, noting that it contains binary gene expression, which can be effectively minimized with the contrastive learning, we optimize the following loss function: 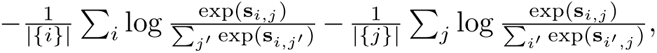 where **s***_i,j_* is the dot product between the paired histopathology image and captions, and has the positive artificial gene expression. The remaining are for non-paired ones with zero artificial gene expression. The summation operators 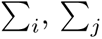 respectively enumerate the image and text indices over the batch, and |{*i*}| and |{*j*}| are respectively the size of histopathology image and captions. Similarly, this applies to 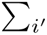 and 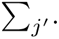 We use the AdamW optimizer with a learning rate of 5 × 10*^−^*^4^, adjusted using a cosine annealing schedule, with a minimum learning rate set to 1 × 10*^−^*^6^. We set the gradient clip norm to 10. During training, we apply a random resized crop with a scale range of 0.9 to 1.0, a horizontal flip with a probability of 0.5, and a vertical flip with a probability of 0.5. The images are then normalized using mean of 0.48145466, 0.4578275, 0.40821073 and standard deviation of 0.26862954, 0.26130258, 0.27577711 for the red, green, and blue channels, respectively.

### Training Platform

The model is trained and evaluated using a system equipped with 8 H100 GPUs, each featuring 96GB of memory, and running driver version 550.54.15. The CPU is an Intel Xeon Platinum 8425Y, complemented by 1024GB of RAM. The implementation is built using Python 3.10.14, PyTorch 2.3.0, CUDA cu121, Torchvision 0.18.0, Einops 0.7.0, Transformers 4.44.0, Regex 2024.4.16, Ftfy 6.2.0, Pandas 2.2.2, Numpy 1.26.3, Webdataset 0.2.86, Braceexpand 0.1.7, Timm 0.9.16, Pillow 9.4.0, and Open clip 2.29.0. Implementations of comparison state-of-the-art methods can be found at OpenAICLIP (https://huggingface.co/openai/ clip-vit-base-patch16), PMCCLIP (https://huggingface.co/ryanyip7777/pmc vit l 14), PathGenCLIP (https://pub-7a38cc906afa44a4a01533c288d0b1af.r2.dev/pathgenclip. pt), BiomedCLIP (https://huggingface.co/microsoft/BiomedCLIP-PubMedBERT256-vit base patch16 224), PLIP (https://huggingface.co/vinid/plip), QuiltNet (https://huggingface.co/wisdomik/QuiltNet-B-32), PathCLIP (https://huggingface.co/jamessyx/pathclip), PubmedCLIP (https://huggingface.co/flaviagiammarino/ pubmed-clip-vit-base-patch32), and CONCH (https://huggingface.co/MahmoodLab/ conch).

### Evaluation

To comprehensively evaluate the potential of AGP-Net, we perform experiments across a range of downstream tasks, including gene expression prediction, tumor detection, tumor and cancer typing, and gene expression imputation. These diverse settings allow us to assess robustness, generalizability, and effectiveness of AGP-Net in addressing complex biomedical challenges. Evaluation details for each task are provided below.

We use PCC (i. e., Pearson correlation coefficient) for evaluations of gene expression prediction and imputation. Following [5], we focus on the 250 most highly expressed genes for evaluations. The textual descriptions of the selected genes are scraped from mygene.info, ensembl.org, and ncbi.nlm.gov. In the gene expression prediction task, the model is tested on out-of-domain spatial transcriptomics datasets. The PCC is calculated between the predicted and ground truth gene expression values for each evaluation dataset.

For the gene expression imputation task, different experimental settings are considered, involving 4, 8, 16, 32, or 64 spatial spots. The corresponding spots are used to crop 224×224 pixel patches from histopathology images, along with their paired gene expression values for training. The remaining spots in each histopathology image are used for evaluation, and PCC is calculated for each histopathology image to assess the model performance. For this task, a linear layer is appended to the image encoder to facilitate gene expression imputation. To address variability in the imputation task, each method is run five times on each slide in the dataset. The model is trained using a batch size of 8 (or 4 for 4-spot settings) with a learning rate of 1 × 10*^−^*^4^, utilizing the AdamW optimizer for 10 epochs.

To evaluate the performance of AGP-Net on tumor detection and tumor and cancer typing tasks, we use accuracy as the primary evaluation metric. Trained on the real and artificial spartial transcriptomics datasets, the model is tested for the classification tasks, with performance measured based on the percentage of correct classifications (i. e., accuracy). To enhance robustness and effectiveness, we construct prompt ensembles following the methodologies described in [16, 30, 44].

## Data Availability

All datasets used in the paper are publicly available. The training and testing real spatial transcriptomics datasets and gene expression imputation are available at STimage-1K4M (https://huggingface.co/datasets/jiawennnn/STimage-1K4M) and HEST-1k (https://huggingface.co/datasets/MahmoodLab/hest). The histopathology image caption datasets used in training are available at PathGen dataset (https://github.com/PathGen-1-6M/PathGen-1.6M), PathCap dataset (https://github.com/superjamessyx/Generative-Foundation-AI-Assistant-for-Pathology, Quilt-1M dataset (https://github.com/wisdomikezogwo/quilt1m), and OpenPath dataset (https://github.com/PathologyFoundation/plip). The dataset for benchmarking tumour detection and tumour and cancer typing are available at LC-Colon dataset (https://academictorrents.com/ details/7a638ed187a6180fd6e464b3666a6ea0499af4af), WSSSLUAD dataset (https://wsss4luad.grand-challenge.org/), SkinCancer-Tumor dataset (https://heidata. uni-heidelberg.de/dataset.xhtml?persistentId=doi:10.11588/data/7QCR8S), NCT-CRC-Tumour dataset (https://zenodo.org/records/1214456), BACH dataset (https://iciar2018-challenge.grand-challenge.org), BreakHis dataset (https://web.inf.ufpr.br/ vri/databases/breast-cancer-histopathological-database-breakhis/), LC-Lung dataset (https://academictorrents.com/details/7a638ed187a6180fd6e464b3666a6ea0499af4af), LungHist700 dataset (https://figshare.com/articles/dataset/LungHist700 A Dataset of Histological Images for Deep Learning in Pulmonary Pathology/25459174), SICAP dataset (https://data.mendeley.com/datasets/9xxm58dvs3/1), and Skin-Cancer dataset (https://heidata.uni-heidelberg.de/dataset.xhtml?persistentId=doi:10.11588/data/7QCR8S).

## Code Availability

Our code can be assessed at TBC.

## Acknowledgments

Liyuan Pan’s work was supported in part by the BIT Special-Zone and National Natural Science Foundation of China 62302045.

## Author Contributions

All author conceived the AGP-Net project and revised the manuscript.

## Competing Financial Interests

The authors declare no competing financial interests.

## Footnotes

1 Broadly speaking, it maps images and text into a shared feature space, allowing universal recognition by measuring their feature similarity.

## Notes

### Competing Interest Statement

The authors have declared no competing interest.

